# DeepImpute: an accurate, fast and scalable deep neural network method to impute single-cell RNA-Seq data

**DOI:** 10.1101/353607

**Authors:** Cedric Arisdakessian, Olivier Poirion, Breck Yunits, Xun Zhu, Lana X. Garmire

## Abstract

**Background:** Single-cell RNA sequencing (scRNA-seq) offers new opportunities to study gene expression of tens of thousands of single cells simultaneously. However, a significant problem of current scRNA-seq data is the large fractions of missing values or “dropouts” in gene counts. Incorrect handling of dropouts may affect downstream bioinformatics analysis. As the number of scRNA-seq datasets grows drastically, it is crucial to have accurate and efficient imputation methods to handle these dropouts.

**Methods:** We present DeepImpute, a deep neural network based imputation algorithm. The architecture of DeepImpute efficiently uses dropout layers and loss functions to learn patterns in the data, allowing for accurate imputation.

**Results:** Overall DeepImpute yields better accuracy than other publicly available scRNA-Seq imputation methods on experimental data, as measured by mean squared error or Pearson’s correlation coefficient. Moreover, its efficient implementation provides significantly higher performance over the other methods as dataset size increases. Additionally, as a machine learning method, DeepImpute allows to use a subset of data to train the model and save even more computing time, without much sacrifice on the prediction accuracy.

**Conclusions:** DeepImpute is an accurate, fast and scalable imputation tool that is suited to handle the ever increasing volume of scRNA-seq data. The package is freely available at https://github.com/lanagarmire/DeepImpute

## Introduction

The RNA sequencing technologies keep evolving and offering new insights to understand biological systems. In particular, single-cell RNA sequencing (scRNA-seq) represents a major breakthrough in this field. It brings a new dimension to RNA-seq studies by zooming in to the single-cell level. Currently, various scRNA-seq platforms are available such as Fluidigm and Drop-Seq based methods. While Drop-Seq can process thousands of cells in a single run, Fluidigm generally processes fewer cells but with a higher coverage. In particular, 10X Genomics’ platform is gaining popularity in the scRNA-seq community due to its high yield and low cost per cell. Consequently, an increasing amount of studies have taken advantage of these technologies to discover new cell types(Usoskin *et al.*, 2015; Villani *et al.*, 2017), new markers for specific cell types(Usoskin *et al.*, 2015; Zeisel *et al.*, 2015; Jaitin *et al.*, 2014), and cellular heterogeneity(Jaitin *et al.*, 2014; Kriegstein *et al.*, 2014; Treutlein *et al.*, 2014; Tirosh *et al.*, 2016; Shalek *et al.*, 2013; Tang *et al.*, 2010).

Despite these advantages, scRNA-seq data are very noisy and incomplete(Kim *et al.*, 2015; Jia *et al.*, 2017; Kolodziejczyk *et al.*, 2015) due to the low starting amount of mRNA copies per cell. Datasets with more than 70% missing (zero) values are frequently observed in an scRNA-seq experiment. These apparent zero values could be truly zeros or false negatives. The latter phenomenon is called “dropout”(Andrews and Hemberg, 2016), and is due to failure of amplification of the original RNA transcripts. Shorter genes are more likely to be dropped out since it is more difficult for RNA polymerase to capture their mRNAs(Pierson and Yau, 2015). Such bias may increase further during the subsequent amplification steps. As a result, dropout can affect downstream bioinformatics analysis significantly, such as clustering(Zhu, Ching, *et al.*, 2017) and pseudo-time reconstruction(Poirion *et al.*, 2017), as it decreases the power of the studies and introduces biases in gene expression. To correct such issue, analysis platforms such as Granatum(Zhu, Wolfgruber, *et al.*, 2017) have included an imputation step, in order to improve the downstream analysis.

Currently several imputation algorithms have been proposed, based on different principles and models. MAGIC(van Dijk *et al.*, 2017) focuses on cell/cell interactions to build a Markov transition matrix and smooth the data. ScImpute(Li and Li, 2017) builds a LASSO regression model for each cell and imputes them iteratively. SAVER(Huang *et al.*, 2017) is a Bayesian-based model using various prior probability functions. DrImpute(Kwak *et al.*, 2017) is a clustering-based method and uses a consensus strategy: it estimates a value with several cluster priors or distance matrices and then imputes by aggregation. As the low quality of the scRNA-seq datasets continues to be a bottleneck while the measurable cell counts keep increasing, the demand for faster and scalable imputation methods also keeps increasing(Eraslan *et al.*, 2018; Lin *et al.*, 2017; Ronen and Akalin, 2018). While some of these earlier algorithms do improve the quality of original datasets and preserve the underlying biological variance(Zhang and Zhang, 2017), often these methods demand extensive running time, impeding their adoption in the ever increasing scRNA-seq data space.

Here, we present a novel algorithm, DeepImpute, for scRNA-seq data imputation. DeepImpute is short for “Deep neural network Imputation”. As reflected by the name, it belongs to the class of deep neural-network models(Ching, Zhu, *et al.*, 2018; Alakwaa *et al.*, 2018; Chaudhary *et al.*, 2018). Recent years, deep learning and related deep neural network algorithms have gained much interest in the biomedical field(Ching, Himmelstein, *et al.*, 2018), ranging from applications from extracting stable gene expression signatures in large sets of public data(Tan *et al.*, 2017) to stratify phenotypes (Beaulieu-Jones *et al.*, 2016) or impute missing values (Beaulieu-Jones and Moore, 2017) using electronic health record (EHR) data. Here, we construct DeepImpute models by splitting the genes into subsets and builds sub-networks to increase its efficacy and efficiency. Using accuracy metrics, we demonstrate that DeepImpute performs better than the four most recent and representative imputation methods mentioned above (MAGIC, DrImpute, ScImpute and SAVER). We also show the superiority of DeepImpute over the other methods in terms of computational running time and memory use. Moreover, DeepImpute allows to train the model with a subset of data to save computing time, with little sacrifice on the prediction accuracy. In summary, DeepImpute is a fast, scalable, and accurate imputation method capable of handling the ever increasing scRNA-seq data.

## Methods

### The workflow of DeepImpute

DeepImpute is an imputation workflow using deep-neural networks implemented with the TensorFlow(Abadi *et al.*, 2016) framework. The algorithm starts by setting a threshold that determines how many genes are to be imputed. This threshold can either be set as the default or determined by the user. As the default, we filter all the genes with less than 5 reads among 99% of the samples. For efficiency, we adopt a divide-and-conquer strategy in our deep learning imputation process. We split the genes into random subsets, each with S numbers of genes, which we call “target genes”. By default S is set as 500.

For each subset, we train a neural network made of four layers: the input layer of genes that are correlated to the target genes, a 300-neuron fully connected hidden layer with a Rectified Linear Unit (ReLU) activation function, a dropout layer (50% drop-out rate), and an output layer made of S “target genes’. A gene is selected into the input layer, if it satisfies: (1) it is not one of the target genes; (2) it has at least 10 reads among 1% of the cells; (3) It has top 20 ranked Pearson’s correlation coefficient with a target gene. The dropout layer is included after the hidden layer, as a common strategy to prevent overfitting(Srivastava *et al.*, 2014). The other default parameters of the networks include a learning rate of 0.0005, a batch size of 64, and a subset size of 500.

In order to emphasize the accuracy on high confidence values during the training phase, we use a weighted MSE for the loss function: *Loss* = Σ *Y*_*i*_·(*Y*_*i*_ − *Ŷ*_*i*_)^2^ where *Y*_*i*_ is the value of gene i, and *Ŷ*_*i*_ is the estimated value for this gene at a given epoch. For the gradient descent algorithm, we choose an adaptive learning rate method, the Adam optimizer(Kingma and Ba, 2014), since it is known to perform very efficiently over sparse data(Ruder, 2016). In addition, we discard cells with poor gene expression values (more than 90% zeros in all cells) during the training.

### scRNA-seq Datasets

In this study, we evaluate imputation metrics on four datasets. Three of them (Jurkat, 293T, neuron9k and Mouse1M) are downloaded from the 10X Genomics support website (https://support.10xgenomics.com/single-cell-gene-expression/datasets). Briefly, the Jurkat dataset is extracted from the Jurkat cell line (human blood). 293T is a blood cell line derived from HEK293T that expresses a mutant version of the SV40 large T antigen. The neuron9k dataset contains brain cells from an E18 mouse. Mouse1M also contains brain cells from an E18 mouse. The fourth data are taken from GEO (GSE67602)(Joost *et al.*, 2016), composed of mouse interfollicular epidermis cells.

### Other methods of comparison

For comparison, we use the latest version of SAVER (v0.3.1) at https://github.com/mohuangx/SAVER/releases, ScImpute (v0.0.6) at https://github.com/Vivianstats/scImpute, DrImpute (v1.0) available as a CRAN package, and MAGIC (pulled 4/3/18) at https://github.com/KrishnaswamyLab/magic. We preprocess the datasets according to each method’s standard: using log transformation for dataset in MAGIC and DeepImpute, but raw counts for scImpute, DrImpute and SAVER.

### Evaluation metrics

#### Speed and memory comparison

We run comparisons on a dedicated 8-core, 30GB RAM, 100GB HDD, Intel Skylake machine running Debian 9.4. We record process memory usage at 60-second intervals. For testing data, we use the Mouse1M dataset since it has the largest number of single cells (**Table 1**). We filter out genes that are expressed in less than 20% of cells, leaving 3205 genes in our sample. From this dataset, we generate 7 subsets ranging in size (100, 500, 1k, 5k, 10k, 30k, 50k cells). We run each package 3 times per subset to estimate the average computation time. Some packages (DrImpute, SAVER, scImpute and MAGIC) are not able to successfully handle the larger files either due to out-of-memory errors (OOM) or exceedingly long run times (> 12 hours).

**Table 1:**
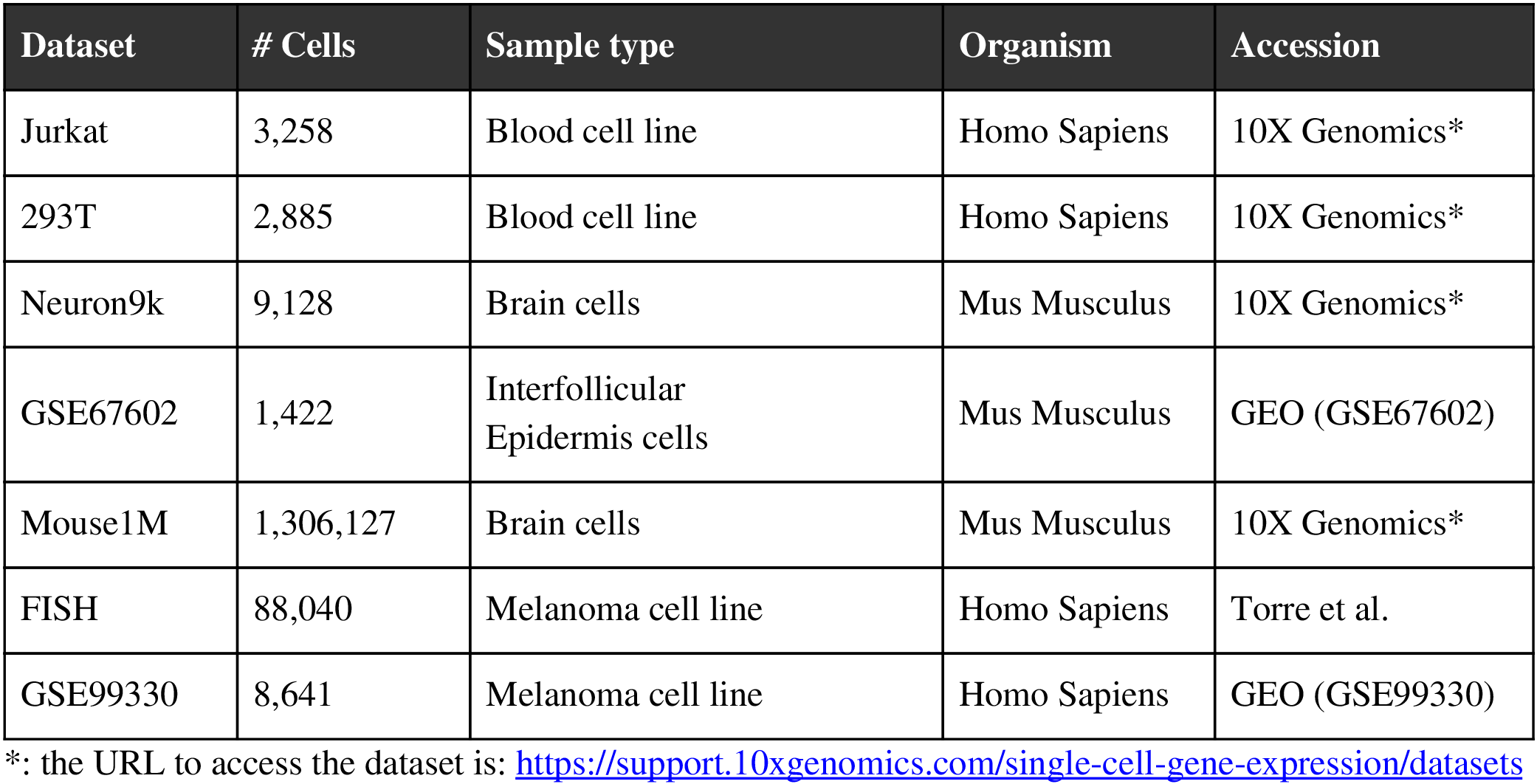
Summary of the single-cell datasets

| Dataset | # Cells | Sample type | Organism | Accession |
| --- | --- | --- | --- | --- |
| Jurkat | 3,258 | Blood cell line | Homo Sapiens | 10X Genomics* |
| 293T | 2,885 | Blood cell line | Homo Sapiens | 10X Genomics* |
| Neuron9k | 9,128 | Brain cells | Mus Musculus | 10X Genomics* |
| GSE67602 | 1,422 | Interfollicular Epidermis cells | Mus Musculus | GEO (GSE67602) |
| Mouse1M | 1,306,127 | Brain cells | Mus Musculus | 10X Genomics* |
| FISH | 88,040 | Melanoma cell line | Homo Sapiens | Torre et al. |
| GSE99330 | 8,641 | Melanoma cell line | Homo Sapiens | GEO (GSE99330) |

\*: the URL to access the dataset is: https://support.10xgenomics.com/single-cell-gene-expression/datasets

#### Accuracy comparison on real datasets

We choose to randomly mask 5% of the positive read count values for each cell in the expression matrix. The masking follows a density probability distribution proportional to *f*(*x*) = *exp*(− *x*/20) where x is the raw read count. These original values are used as the reference to evaluate the performance of imputation methods. We used two types of performances metrics: the overall Pearson’s correlation coefficient and the mean squared error (MSE), both on log transformed counts. When needed, we also computed MSE between cells *C*_*j*_ and between genes *g*_*i*_.

### RNA FISH validation

We obtain a Drop-Seq dataset (GSE99330) and its RNA FISH dataset from a melanoma cell line, as described by Torre et al(Torre *et al.*, 2018). The summary of the dataset is listed in **Table 1**. For the comparison between RNA FISH and the corresponding Drop-Seq experiment, we keep the top 2000 genes as done on other datasets in this study that are used for evaluations, leaving six genes in common between the FISH and the Drop-Seq datasets. We normalized the cells in each dataset using a housekeeping gene (Glyceraldehyde 3-phosphate Dehydrogenase, or GAPDH) based factor: we multiply each cell by its mean(GAPDH)/GAPDH value. To compute Gini index, we filter genes below the 10^th^ and above the 90^th^ percentile, as done by others(Huang *et al.*, 2017).

### Software and code availability

DeepImpute package and its documentation are freely available at https://github.com/lanagarmire/DeepImpute.

## Results

### Overview of the DeepImpute algorithm

DeepImpute is a deep neural network model that imputes genes in a divide-and-conquer approach, by constructing multiple sub-neural networks. In each sub-neural network, it aims to understand gene networks by splitting them between the input and the output layer. Doing so offers the advantage of reducing the complexity by learning smaller problems and fine-tuning the sub-neural networks(Chiang and Fu, 1994). Users can set the size of the subset data. We use 500 genes as the default value, since it offers a good trade-off between speed and stability. For each sub-model, we train a feed-forward neural network to fit these 500 genes, which we call target genes.

As shown in **Figure 1**, each sub-neural network is composed of four layers. The input layer consisting of genes that are correlated with the target genes and have read counts above 10 in at least 1% among all cells. It is followed by a 300-neuron dense hidden layer, a dropout layer with 50% drop-out rate of neurons (to avoid overfitting), and the output neurons made of the above mentioned target genes. We use Rectified Linear Unit (ReLU) as the activation function, and train each sub-model using data from cells with at least 10% non-zero values. Because of the simplicity of each sub-network, we observe very low variability due to hyperparameter tuning. As a result, we set the default parameters for batch size at 64 and learning rate at 0.005. Further information about the network parameters are described in **Methods**. In the following sections, we performed comprehensive evaluations of DeepImpute by combining in parallel the results of all sub-networks.

**Figure 1:**
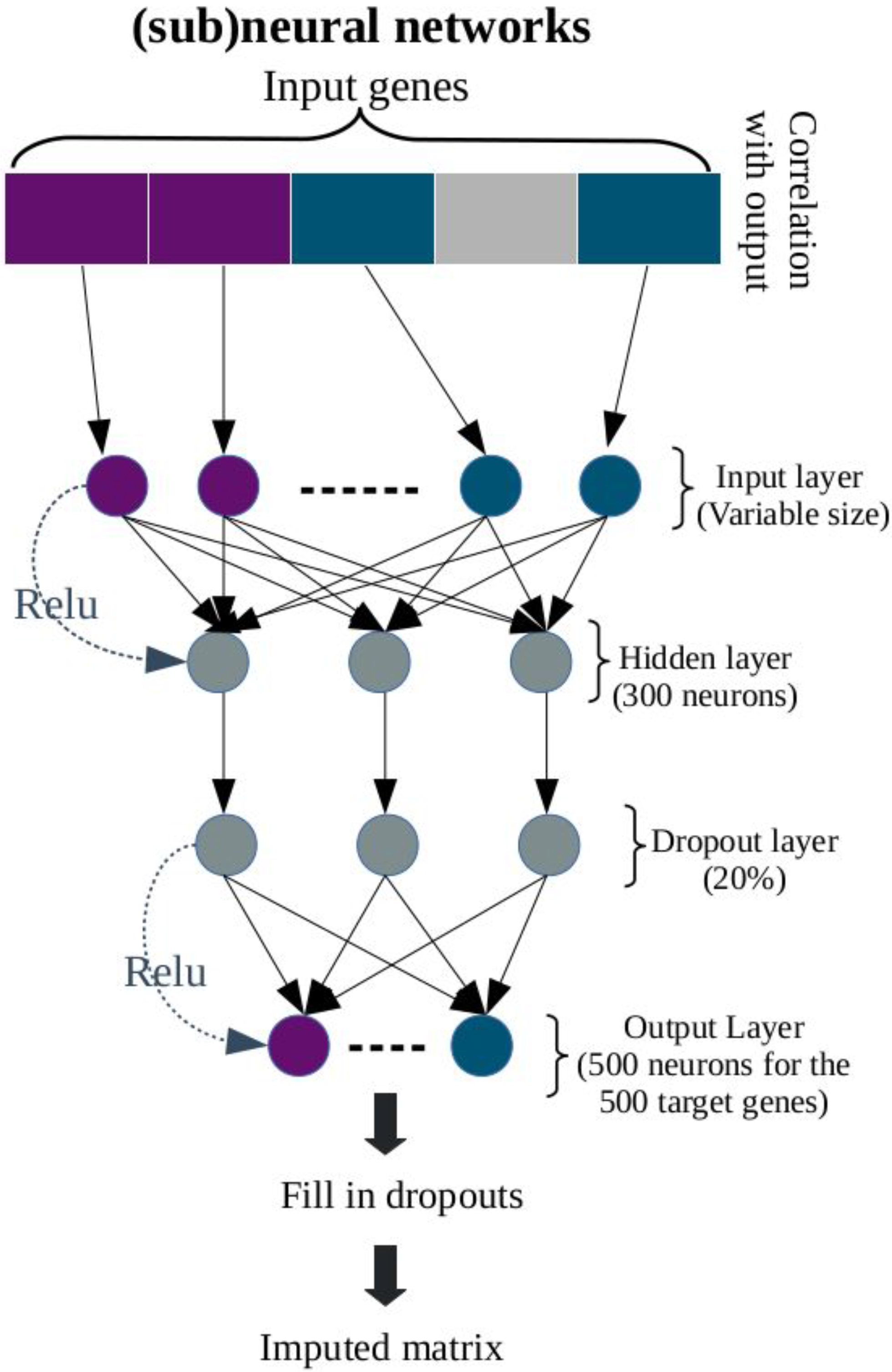
(Sub) Neural network architecture of DeepImpute. For each sub neural network, the input layer consists of genes that are correlated with the genes in the output layer. It is followed by two hidden layers: one 300 neurons dense layer and a dropout layer (rate=50%). The output layer consists of the subset of genes randomly assigned (default N=500).

### DeepImpute is the most accurate among imputation methods on scRNA-seq data

We tested the results on four publically available scRNA-seq datasets (**Table 1**): two cell lines, Jurkat and 293T (10X Genomic), one mouse neuron cells dataset (10X Genomics), and one mouse interfollicular epidermis (IFE) dataset deposited in GSE67602. We compared DeepImpute with four other state-of-the-art, representative algorithms: MAGIC, DrImpute, ScImpute and SAVER. Since the real dropout values are unknown, we evaluated the different methods by randomly masking (replacing with zeros) a part of the expression matrix of a scRNA-seq dataset, and then measure the differences between the inferred and actual values of the masked data. In order to mimic a more realistic dropout distribution, we masked the lower values with a higher probability following the exponential law (see **Methods**). To keep a reasonable number of genes for imputation, we studied the performances on the top 2000 genes chosen by the 99% quantiles’ count values. This threshold allows to obtain the majority of genes that have more than 5 reads in 1 percentile of cells.

We measured the accuracies using the two metrics on the masked values: Pearson’s correlation coefficient and Mean Squared Error (MSE), as done earlier(Garmire and Subramaniam, 2012; Huang *et al.*, 2017). **Figure 2** shows all the results of imputation accuracy metrics on the masked data. DeepImpute successfully recovers dropout values from all ranges, introduces the least distortions and biases to the masked values and yields both the highest Pearson’s correlation coefficient and the best (lowest) MSE in each dataset (**Figure 2A**). On the contrary, other methods present various issues: MAGIC has the 2nd best Pearson’s correlation coefficients after DeepImpute, but worse MSEs overall compared to SAVER. SAVER presents systematic bias to underestimate the masked data, especially in the higher value range, thus it produces overall worse Pearson’s correlation coefficients than MAGIC. DrImpute and scImpute have the widest ranges of variations among imputed data, compared to those of masked values, therefore giving two worst sets of Pearson’s correlation values. The variation of imputed data from scImpute is the largest, and it produces the worst (highest) overall MSE. We further examined MSE distributions calculated on the gene and cell levels (**Figure 2B and 2C**). DeepImpute is the clear winner with consistently the best (lowest) MSEs for gene and cell levels, on all datasets. scImpute consistently gives the worst (highest) MSEs at the cell level and 2nd worst MSEs at the gene level (Figure 2B and 2C). Other methods are ranked in between, with varying rankings depending on the datasets and gene or cell level calculations. In summary, DeepImpute yields the highest accuracy in the datasets studied, among the imputation methods in comparison.

**Figure 2:**
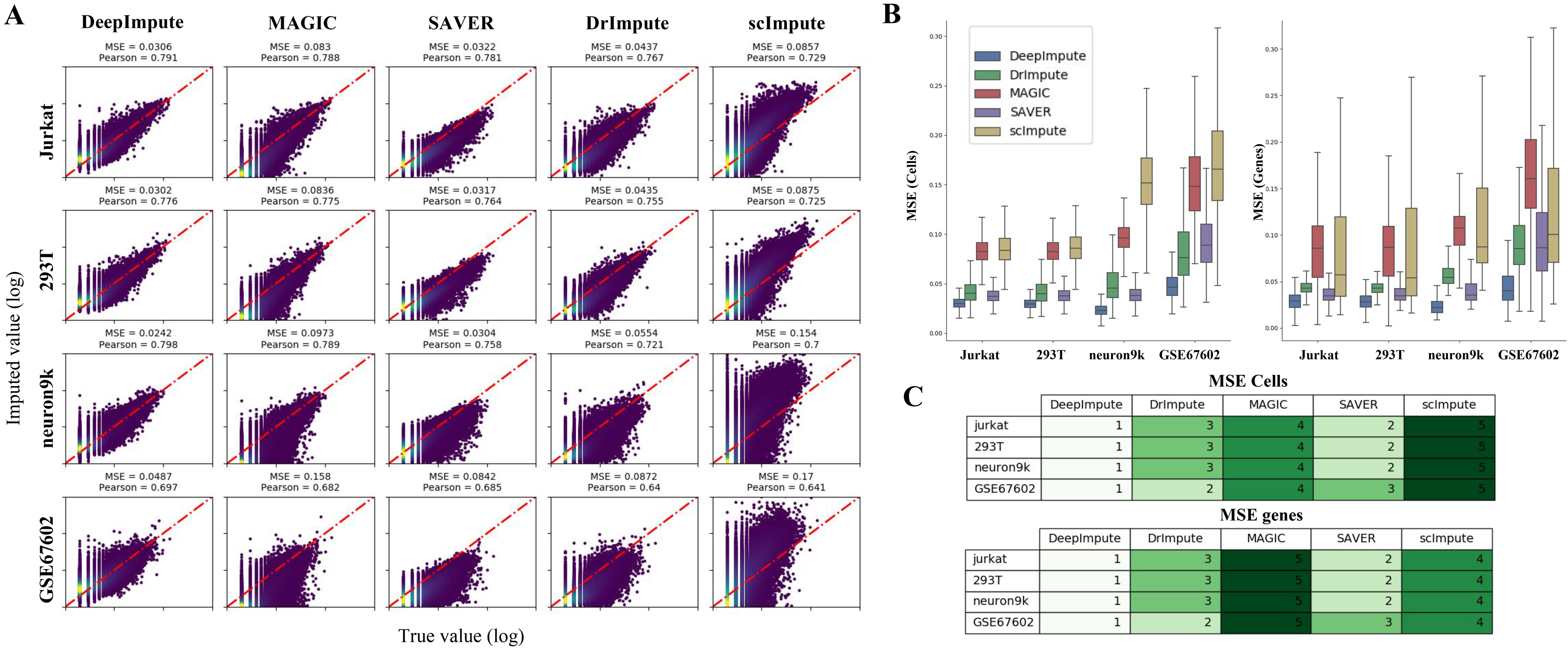
Accuracy comparison between DeepImpute and other competing methods. (**A**) Scatter plots of imputed vs. original data masked. The x-axis corresponds to the true values of the masked data points, and the y-axis represents the imputed values. Each row is a different dataset, and each column is a different imputation method. The mean squared error (MSE) and Pearson’s correlation coefficients (Pearson) are shown above each dataset and method. (**B**) Bar graphs of cell-cell and gene-gene level MSEs between the true (masked) and imputed values, based on those in (**A**). Color labels: DeepImpute (blue), DrImpute (green), MAGIC (red), SAVER (purple) and scImpute (gold) on real datasets. (**C**) Heatmap rankings of MSEs, for the conditions in (**B**). Lighter vs. darker green indicates better vs. worse rankings.

### DeepImpute improves the gene distribution similarity with FISH experimental data

Another way to assess the imputation efficiency is through experimental validation on scRNA-Seq data. Single-cell RNA FISH is such a method that directly detects a small number of RNA transcripts in a single cell. Torre *et al.* measured the gene expression of a melanoma cell line using both RNA FISH and Drop-Seq and compared their distribution using their Gini coefficients (see **Methods**)(Torre *et al.*, 2018). Similarly, we compared the same list of genes using their Gini coefficients of RNA FISH vs. those after imputation (or raw the scRNA-seq data). Comparing to the Pearson’s correlation coefficient between RNA FISH and the raw scRNA-seq data (−0.369), three methods, DeepImpute, DrImpute and SAVER, have improved correlation coefficients. Among them, DeepImpute had the highest value (0.944). On other other hand, MAGIC and scImpute both had worse (lower) correlation coefficients than the raw scRNA-seq dataset (**Figure 3**). For MSE, all imputation methods achieved better (smaller) MSEs compared to the raw scRNA-seq results (MSE=0.281). Still, DeepImpute is the most accurate method with an MSE (MSE=0.0204). This MSE is also much less compared to the second lowest MSE (MSE=0.0437) generated from DrImpute. Thus, these results demonstrate that DeepImpute improves the data quality the best among all compared methods, using RNA FISH experimental data as the truth measure.

**Figure 3:**
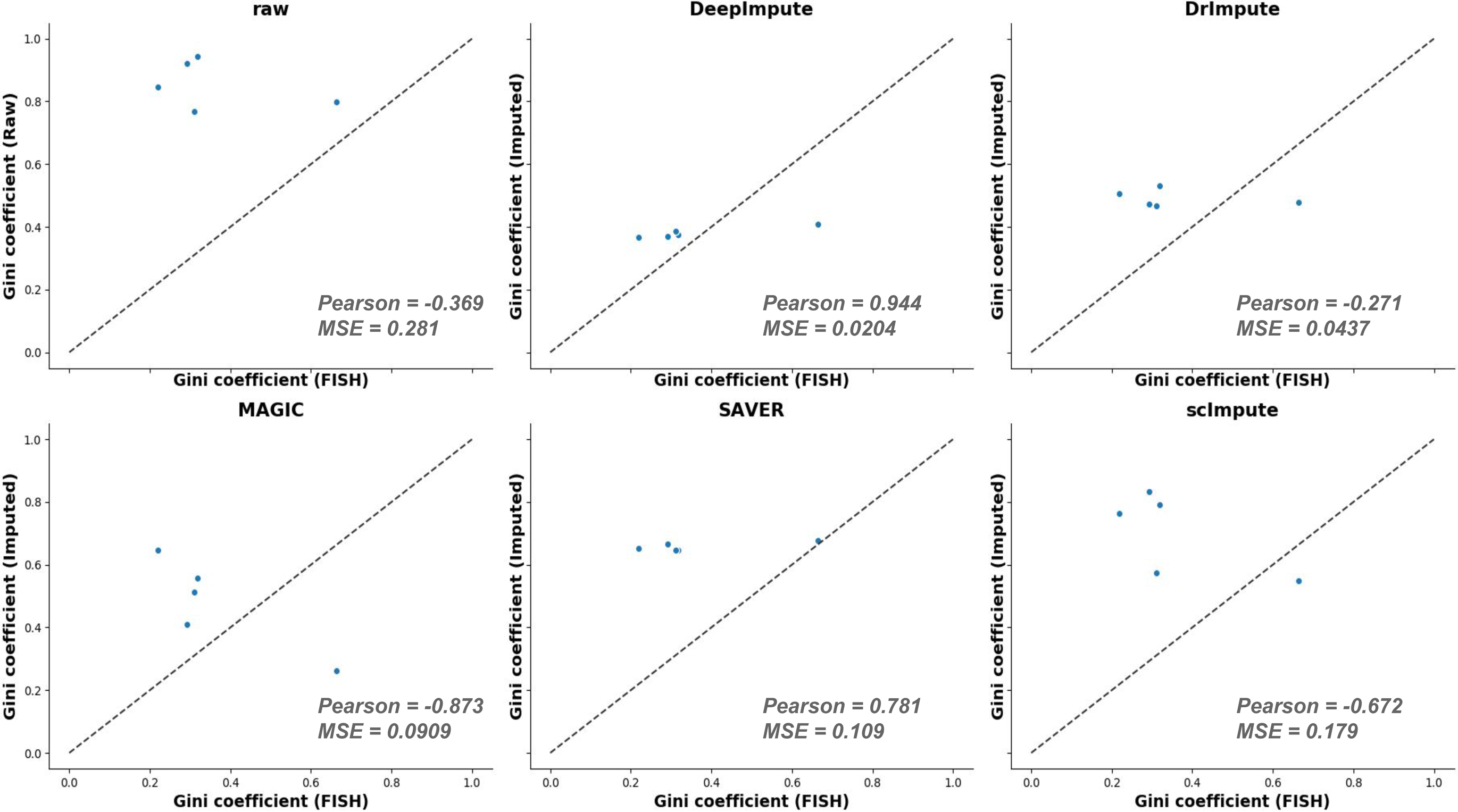
Comparison among imputation methods using RNA FISH data. (**A**) Scatter plots of Gini index from the imputed (or raw) vs. FISH data. The x-axis is the “true” Gini coefficient as determined by FISH experiments, and the y-axis is the imputed (or raw) Gini coefficient. The Pearson’s correlation coefficients (Pearson) and mean squared error (MSE) are shown for each method.

### DeepImpute is a fast and memory efficient package

As scRNA-seq becomes more popular and the number of sequenced cells scales exponentially, imputation methods will have to be computationally efficient to be widely adopted. With such a goal in mind, we chose the Mouse1M dataset to evaluate the computational speed and memory usage among different imputation methods. We used Mouse1M dataset as it has the highest number of cells to assess how adaptive each method is.

We downsampled the data 7 subsets ranging in size from 100 to 50k cells (100, 500, 1k, 5k, 10k, 30k, 50k). We ran the imputations three times and measured the runtime and memory load on an 8-core machine with 30 gigabytes of memory. Overall, DeepImpute and MAGIC outperformed the other three packages on both speed (**Figure 4A**) and memory usage, especially on large datasets (**Figure 4B**). DeepImpute and MAGIC performed similarly on smaller datasets, however, as the dataset size increased their memory usage diverged (**Figure 4B**). On the 30k cell dataset, DeepImpute consumed less than half (~8GB) of the available memory, while MAGIC used approximately ⅔ of the memory (~20GB). On the 50K cell dataset, MAGIC hit an out of memory error and was unable to finish the 50K cell imputation on our 30GB machine. The other three imputation methods (scImpute, DrImpute and SAVER) were significantly slower and consumed significantly more memory (**Figure 4**). The slow computation time of DrImpute is due to lack of parallelization. Furthermore, both SAVER and scImpute exceeded the 30GB of memory available and failed to run on more than 10k cells. Therefore, judging by both computation speed and memory efficiency on larger datasets, DeepImpute again tops the other methods.

**Figure 4:**
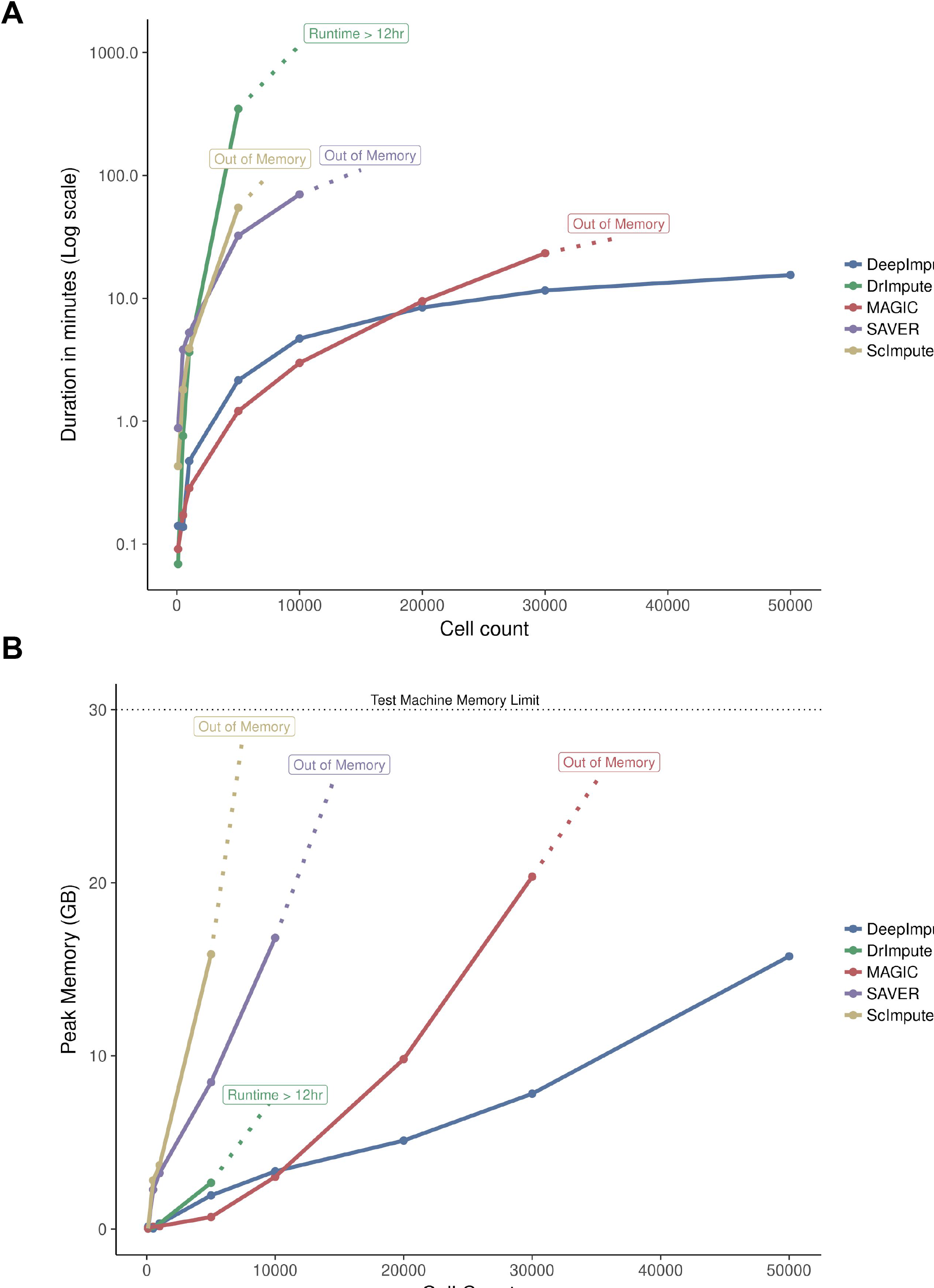
Speed and memory usage comparison among imputation methods, on the Mouse1M dataset. This dataset is chosen for its largest cell numbers. Color labels: DeepImpute (blue), DrImpute (green), MAGIC (red), SAVER (purple) and scImpute (gold). (A) Speed evolution over 3 runs. The x-axis is the number of cells and the y-axis is the running time in minutes (log scale) of the imputation process. (B) RAM memory usage. The x-axis is the number of cells and the y-axis is the maximum RAM used by the imputation process. Because of the limited amount of memory or time, scImpute, SAVER and MAGIC exceeded the memory limit respectively at 10k, 30k and 50k cells. DrImpute exceeded 12h on 10k.

### DeepImpute is a scalable machine learning method

Unlike the other imputation methods, DeepImpute first fits a predictive model and then performs imputation separately. The model fitting step uses most of the computational resources and time, while the prediction step is very fast. We then asked the question what is the minimal fraction of the dataset needed to train DeepImpute and obtain efficient imputation without extensive training time. Hence, we used the neuron9k dataset and evaluated the effect of different subsampling fraction (5%, 10%, 20%, 40%, 80%, 100%) in the training phase on the imputation prediction phase. We randomly picked a subset of the samples for the training and computed the accuracy metrics (MSE, Pearson’s correlation coefficient) on the whole dataset, with 10 repetitions under each condition. Model performance improvement begins to slow down at around 40% of the cells (**Figure 5**). Specifically, from 40% to 100% fraction of data as the training set, the MSE decreases slightly from 3.69 to 3.43, and Pearson’s coefficient score marginally improves from 0.863 to 0.875. These experiments demonstrate another advantage of DeepImpute over the competing methods, that is, the use of only a fraction of the data set will reduce the running time even more with little sacrifice to the accuracy of the imputed results.

**Figure 5:**
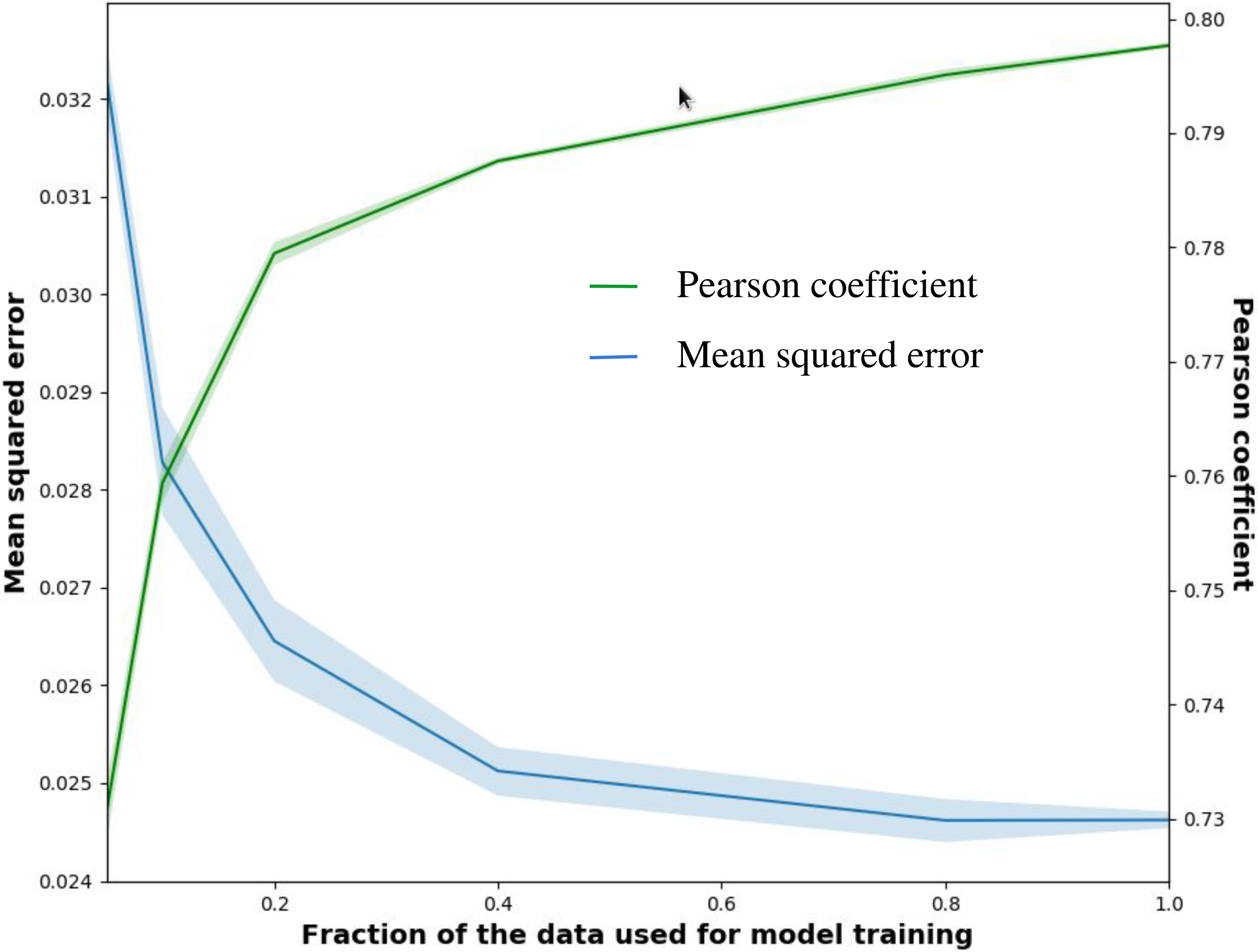
Effect of subsampling the training data on imputation accuracy. neuron9k dataset is masked and measured for performance as in Figure 2. X-axis is the fraction of cells in the training data set, and y-axis are values for mean squared error (left) or Pearson’s correlation coefficient (right). Color label: blue line (mean squared error), green line (Pearson’s correlation coefficients). Shades represent the variations over the 10 repetitions.

## Discussion

Dropout values in scRNA-seq experiments represent a serious issue for bioinformatic analyses, as most bioinformatics tools have difficulty handling sparse matrices. In this study, we present DeepImpute, a new algorithm that uses deep neural networks to impute dropout values in scRNA-seq data. We show that DeepImpute not only has the highest overall accuracy, but also offers faster computation time with less demand on the computer memory. Furthermore, it is a very “resilient” method. The model trained on a fraction of the input data can still yield decent predictions, which can further reduce the running time. Together, these results demonstrate that DeepImpute is an accurate and highly efficient method, and it is likely to withstand the tests of time, given the rapid growth of scRNA-Seq data volume.

Several unique properties of DeepImpute contribute to its superior performance. One of them is using a divide-and-conquer approach. This approach has several benefits. First, contrary to auto-encoders, the subnetworks are trained without using the target genes as the input. It reduces overfitting while enforcing the network to understand true relationships between genes. Second, splitting the genes into subsets results in a lower complexity in each sub-model and stabilizing neural networks. As a result, a small change in the hyperparameters has little effect on the result. Using a single set of hyperparameters, DeepImpute achieves the highest accuracies in all four datasets. Third, splitting the training into sub-networks results in increased speed as there are fewer input variables in each subnetwork. Also, training of each sub-network is done in parellel on different threads, which is more difficult to do with one single neural network.

Unlike other imputation algorithms in comparison, DeepImpute is a machine learning method. The training and the prediction processes of DeepImpute are separate, and this may provide more flexibility when handling large datasets. Moreover, we have shown that using only a fraction of the overall samples, one can still obtain decent imputation results without sacrificing the accuracy of the model much, thus further reducing the running time. Perhaps another advantage of DeepImpute over other methods, is that one can pre-train a dataset of a cell type (or cell state) on another cell type (or cell state) decently. This pre-training process is very valuable in some cases, such as when the number of cells in the dataset is too small to construct a high-quality model. Pre-training can also largely reduce the overall computation time, since DeepImpute spends most of the time on training the samples. Thus it is also a good strategy when the new, large dataset is very similar to the dataset used in pre-training. None of the other competing methods in this study offers such flexibility and time-saving customization.

An enduring imputation method has to adapt to the ever-increasing volume of scRNA-seq data. DeepImpute is such a method, implemented in deep learning framework where new solutions for speed improvement keep appearing. One example is the development of neural networks specific hardware (such as Tensor Processing Units(Jouppi *et al.*, 2017), or TPUs) which are now available on Google Cloud. TPU can dramatically accelerate the tensor operations and thus the imputation process. We were already able to deploy DeepImpute in a Google Cloud environment where TPUs are already available. Another example is the development of frameworks that efficiently use computer clusters to parallelize tasks such as Apache-Spark(Shanahan and Dai, 2017) or Dask(Mehta *et al.*, 2017). Such resources will help DeepImpute and similar methods achieve even higher imputation speed over time, and keep up with the development of scRNA-seq technologies.

## Author Contributions

LG envisioned this project. OP helped supervising CA with LG. CA and BY implemented the project and conducted the analysis with the help from OP, BY and XZ. CA, OP, BY and LG wrote the manuscript. All authors have read and agreed on the manuscript.

## Acknowledgements

The authors thank Dr. Arjun Raj and Eduardo Torre for providing the data for RNA FISH and Drop-seq. This research was supported by grants K01ES025434 awarded by NIEHS through funds provided by the trans-NIH Big Data to Knowledge (BD2K) initiative (https://www.bd2k.nih.gov
), P20 COBRE GM103457 awarded by NIH/NIGMS, R01 LM012373 awarded by NLM, R01 HD084633 awarded by NICHD to L.X. Garmire.

## Supplementary Figures

**Supplementary Figure 1:** 99^th^ percentile distribution across datasets. The x-axis is the sorted gene indices based on their 99^th^ percentile, and the y-axis is the read count value at the 99^th^ percentile. The shaded region corresponds to the genes that have been kept for the comparison, and the count value for the lower limit of each dataset appears on the y-axis.

